# STAT4-dependent regulation of neuroinflammation in atherosclerosis

**DOI:** 10.64898/2026.03.20.713185

**Authors:** Natalie Stahr, Alina K. Moriarty, Shelby D. Ma, W. Coles Keeter, Woong-Ki Kim, Larry D. Sanford, Elena V. Galkina

## Abstract

Atherosclerosis is linked to an increased risk of cognitive decline, with chronic inflammation being a common feature of both pathologies. IL-12 activates STAT4 to regulate myeloid cell functions, and blockade of this pathway alleviates cognitive impairment in Alzheimer’s models. However, the mechanisms connecting vascular pathology to neuroinflammation remain unclear. Here, we examine whether STAT4 functions as a common mediator of neuroinflammation in atherosclerosis.

We demonstrate that LysM^Cre^-specific STAT4 deficiency ameliorates deficits in long-term memory in low-density lipoprotein-deficient (*Ldlr^−/−^*) mice fed a high-fat diet (HFD-C). STAT4 deficiency moderately reduces Ser199-phosphorylated Tau burden. Atherosclerosis alters brain immune composition, characterized by increased numbers of CD45^+^ leukocytes, activated microglia, and activated T and B cells, whereas STAT4 deficiency attenuates these effects. Nanostring gene-expression pathway analysis further highlights the importance of STAT4 in regulating multiple neuroinflammatory pathways and the Rhodopsin-like receptor signaling, which is associated with synaptic plasticity. LysM^Cre^-specific STAT4 deficiency supports microglial efferocytosis in atherosclerotic *Ldlr^−/−^* mice and increases the number of efferocytotic macrophages. Accordingly, STAT4 deficiency also reduced neuronal death.

Overall, our data reveal an important role for myeloid-driven STAT4 expression in the pathogenesis of cognitive decline associated with atherosclerosis, mediated through impaired efferocytosis and enhanced leukocyte activation, leading to increased brain neuroinflammation.

## 1. Introduction

Atherosclerosis is a progressive, chronic inflammatory disorder of the vascular system, characterized by the accumulation of modified lipoproteins, infiltration of immune cells, and the formation of fibrotic tissue within the arterial walls (Keeter et al., 2022; Libby, 2021; Stahr & Galkina, 2022). This disease affects blood vessels throughout the body and can disrupt the function of numerous organs, including the brain (Keeter et al., 2022; Stahr & Galkina, 2022). Atherosclerosis increases the risk of stroke, but has additionally been implicated as a risk factor for cognitive impairment associated with neuroinflammation (Hayes et al., 2025; Stahr & Galkina, 2022). Neuroinflammation is currently considered a driving force in the progression and likely etiology of numerous neurological diseases, including neurodegenerative diseases such as Alzheimer’s disease (AD). A meta-analysis showed that carotid intima-media thickness, a measure of atherosclerosis, is significantly higher in individuals with AD compared to age-matched controls (Xiang, 2017; Xie et al., 2020). Intracranial atherosclerosis is also associated with an increased risk for dementia (Dolan et al., 2010). Furthermore, advanced atherosclerosis can lead to hypoxia in the brain, which contributes to the deposition of amyloid β (Aβ) (Gupta & Iadecola, 2015). Given the growing evidence suggesting the systemic inflammation associated with atherosclerosis triggers neuroinflammation, it is critical to investigate how atherosclerosis affects neuroinflammation and cognitive function.

In preclinical models of AD, a high-fat diet is often used to induce atherosclerosis in mice, leading to increased Aβ deposition, microglial activation, and greater memory impairment (Amelianchik et al., 2022; Bracko et al., 2020; Lin et al., 2016; Mazzei et al., 2021; Nam et al., 2017; Walker et al., 2017). AD is also characterized by a significant efflux of myeloid cells to the brain that are found in areas with Aβ deposits (Zenaro et al.). Common models of atherosclerosis have demonstrated cognitive decline and other characteristics of neurodegeneration. Apolipoprotein E-deficient (*Apoe*^−/−^) mice, for example, exhibit reduced Aβ clearance, increased tau hyperphosphorylation, and impaired performance in the Morris water maze test (Bink et al., 2013). Low-density lipoprotein receptor-deficient *(Ldlr^−/−^)* mice exhibit similar impairments in spatial and working memory in the Morris water maze, object location tasks, and the water radial-arm maze. Importantly, brains from both *Apoe^−/−^* and *Ldlr^−/−^* mice display increased microglial activation (Bink et al., 2013). While several studies have demonstrated associations between atherosclerosis, neuroinflammation, and cognitive impairment, the precise mechanistic pathways connecting vascular pathology to central nervous system (CNS) dysfunction remain unclear.

Interleukin (IL)-12, a pro-inflammatory cytokine, with a main function to induce interferon-gamma (IFNγ) production and promote the differentiation of T helper 1 (Th1) cells (Ullrich et al., 2020). IL-12 is upregulated in the brains of APP/PS1 mice, a commonly used mouse model of AD, and the inhibition of the p40/IL-12 pathway ameliorates amyloid burden and improves cognitive performance in a Barnes maze test in these animals (Vom Berg et al., 2012). New data indicate that IL-12 signaling contributes to AD pathology by disrupting neuronal and oligodendrocyte homeostasis (Schneeberger et al.).

Signal transducer and activator of transcription 4 (STAT4) is the primary transcription factor directly activated following IL-12R signaling (Leonard & O’Shea, 1998). Although traditionally associated with Th1 cell differentiation, IL-12/STAT4 signaling also plays a critical role in neutrophil functions (Mehrpouya-Bahrami et al., 2021) and monocyte differentiation into a pro-inflammatory macrophage (MΦ) phenotype (Fukao et al., 2001; Taghavie-Moghadam et al., 2015). Importantly, myeloid-specific STAT4 deficiency reduces atherosclerotic lesion burden and improves plaque-stability phenotype (Keeter et al., 2023). Neuroinflammation is characterized by significant alterations in the composition and activation of immune cells within the brain. To date, little is known about how atherosclerosis regulates activation at this immune-privileged site. Herein, we examine the role of the IL-12/STAT4 axis in mediating inflammation and cognitive decline in a mouse model of atherosclerosis.

## 2. Materials and Methods

### Animal and Study Design

*Stat4^fl/fl^* mice were generated previously (Mehrpouya-Bahrami et al., 2021) and crossed with *Ldlr^−/−^* mice (JAX, Cat:#002207, RRID:IMSR_JAX:002207) and *B6.129P2-Lyz2tm1(cre)Ifo/J* transgenic mice (JAX, Cat: #004781, RRID:IMSR_JAX:004781) to generate myeloid-specific *Stat4^ΔLysM^Ldlr^−/−^* mice (Keeter et al., 2023). *Stat4^fl/fl^Ldlr^−/−^* mice were used as controls. To induce chronic hypercholesterolemia and advanced atherosclerosis, 8-12-week-old male and female mice were placed on a high-fat diet with 0.15% added cholesterol (HFD-C, fat: 60% kcal; carbohydrate: 26% kcal, Bio-Serv F3282) for 28 weeks until sacrifice. Additionally, 8-12-week-old *Stat4^fl/fl^Ldlr^−/−^* mice were maintained on a chow diet (fat: 13.6% kcal; carbohydrate: 57.5% kcal; LabDiet 5001) for 28 weeks prior to sacrifice, serving as controls. Animals were kept on a 12-hour light/dark cycle with ad libitum access to food and water under pathogen-free conditions. Animal protocols were approved by the Eastern Virginia School Institutional Animal Care and Use Committee and were conducted in accordance with the National Institutes of Health Guidelines for Laboratory Animal Care and Use.

### Behavioral Studies

The Open Field and Novel Object Recognition (NOR) Tests were conducted with HFD-C-fed and chow diet-fed mice during the week preceding sacrifice. Behavioral tests were recorded live and evaluated using Noldus Ethovision (RRID:SCR_000441) automated scoring.

#### Novel Object Recognition

Following the Open Field test, which allowed for acclimation to the chamber, we performed the NOR test (Lueptow, 2017). Two identical objects (caps from 15 mL conical tubes) were placed in opposing corners of the Open Field chamber, 3 inches inwards from the corners. Mice were individually allowed to explore for 8 minutes before being returned to their respective home cages. Twenty-four hours later, the mice were placed in the Open Field chamber and allowed to acclimate for an additional 8 minutes. Subsequently, one familiar object (the cap from the conical tube) was placed in one corner, as before, and a novel object (a 1.5 mL microcentrifuge tube) was placed in the opposite corner. Mice were allowed to explore for 8 minutes, then returned to their respective home cages. Mice that remember the familiar object from the previous day will choose to investigate the novel object for a disproportionately longer time compared to how long they explore the familiar object on the second day. This is quantified by calculating the time spent exploring the novel object minus the time spent exploring the familiar object, then dividing the result by the total exploration time and multiplying by 100. The chamber and objects were cleaned with 70% ethanol before testing and between animals to eliminate odor-related bias from previous animals. Exploration was defined as the mouse’s time spent in proximity to the object (within 2-3 cm), with its head facing the object (Lueptow, 2017).

#### Open Field

The open field test was performed to assess exploratory and anxiety-like behavior (Tang & Sanford, 2005). Mice were placed individually in the center of a 16-inch × 16-inch open-field chamber and allowed to explore freely for 8 minutes. Anxiety-like behavior was determined by calculating the percentage of time spent in the center of the chamber (defined as a centered 8-inch square within the chamber) out of the total time spent in the chamber (Tang & Sanford, 2005). Anxious mice are more likely to avoid the open area at the center of the chamber and stay in the periphery, while less anxious mice will explore the center more willingly. The chamber was cleaned with 70% ethanol before testing and between animals to eliminate bias due to odors of previous animals.

### Tissue collection

Mice were euthanized via CO_2_ asphyxiation. Blood was collected by cardiac puncture, and the heart was perfused with cold PBS to clear blood from the brain vasculature. The whole brain was collected and mechanically dissociated in 1mL PBS using a Dounce homogenizer on ice. The homogenized tissue was sieved through a 70µm cell strainer (Fisher Scientific, cat#22363548) into a 15mL Falcon tube. The homogenizer was washed with PBS, and this PBS was then passed through another 70 µm cell strainer. Homogenates were pelleted at 400×g for 6 minutes at 4°C. The supernatant was removed, and the pellet was resuspended in 500 µL of isotonic Percoll (1.1g/mL, Sigma-Aldrich, cat#P4937) and 3 mL of PBS. 2 ml of PBS were then layered on top. Samples were centrifuged for 10 minutes at 3000xg. Following removal of the upper layers, the cell pellet was washed with 8 mL PBS and centrifuged at 400xg for 10 minutes at 4°C. After aspiration of the supernatant, the pellet was resuspended in cold PBS, and the cells were counted using a hemocytometer. Trypan Blue staining was used to distinguish live and dead cells.

### Flow Cytometry

Brain cells in a single cell suspension were incubated with 1 μL Fc block (1uL per 2×10^6^ cells) for 10 minutes at room temperature. Viability dye was then added and incubated for 10 minutes at room temperature, followed by the addition of the remaining master mix of fluoroconjugated antibodies and an additional 10 minutes of incubation at room temperature. Cells were fixed in 1% formalin for 20 minutes, washed with PBS, and resuspended in PBS before storage at 4°C. If samples required intracellular staining, they were fixed and permeabilized using the True Nuclear Transcription Factor Buffer set (BioLegend, cat# 424401) according to the manufacturer’s instructions. ZoombiNir staining was used to identify apoptotic/dead cells (Biolegend; Cat#423105). The samples were acquired using a 4-laser Cytek® Aurora (Cytek, Inc., RRID:SCR_019826), unmixed using SpectroFlo, and analyzed with FlowJo (Tree Star Inc., RRID:SCR_008520). For all experiments, the gates were set based on the isotype and a fluorescent-minus-one control. Antibodies against CD11b (eFluor450, Cat.#48-0112-82), B220 (eFluor506, Cat.#RA36B2); Tmem119 (PE-Cy7, Cat.#25-6119-82); and CD4 (BV65, Cat.#0416-0042-80) were from ThermoFisher; CD45 (Per-CP, Cat.#103129), CD64 (BV650, Cat.#139337); MerTK (BV711, Cat.#151515); I-Ab (BV7851, Cat.#07645); CD206 (FITC, Cat.#141703); Ly6G (Per-CP-Cy5.5, Cat.#127615), CD3 (PE-Dazzle 594, Cat.#100347); CD69 (SparkNIR; Cat.#104557); Ly6C (AF700, Cat.#128023); CD68 (PE, Cat.#127013); CD11b PB (PB, Cat.#101223); CD19 (APC-Cy7, Cat.#152411); and ZombieNIR Fixable Viability Kit (Cat.#423105) from Biolegend. Antibodies against CD115 (BV480, Cat.#AFS98); CD64 (BV786, Cat.#:BDB569507) were from BD Bioscience, and NeuN (AlexaFluor488, Ab190195) from Abcam.

Microglia were defined as CD45^low/+^CD11b^low^Tmem119^+^ cells; B cells were identified as CD45^+^Tmem119^-^Ly6G^-^CD3^-^CD19^+^ or CD45^+^Tmem119^-^Ly6G^-^CD3^-^B220^+^ cells; T cells were identified as CD45^+^Tmem119^-^Ly6G^-^CD19^-^CD3^+^ cells, and monocytes and macrophages were identified as CD45^+^Tmem119^-^Ly6G^-^CD19^-^CD3^-^ CD11b^+^ cells. Neurons were identified as CD45^-^CD11b^-^NeuN^+^ cells.

### RNA Isolation and Quantitative PCR

Mouse brains were gently homogenized in TRIzol Reagent (Invitrogen, Cat. #15596026) using a Dounce tissue grinder, then chloroform was added and the mixture was combined by inversion. Samples were incubated at room temperature for 10 minutes, then centrifuged at 16,000×g for 5 minutes at 4°C. The aqueous phase was collected and combined with 1 volume of isopropyl alcohol. RNA was then isolated from these samples using the RNeasy Kit for RNA Purification (Qiagen, Cat. # 74104) according to the manufacturer’s instructions. Purity and concentration of RNA were verified using the Nanodrop ND-1000 Spectrophotometer (RRID: SCR_016517). RNA was reverse-transcribed into cDNA using the iScript cDNA Synthesis Kit (Bio-Rad Cat. #74104) according to the manufacturer’s instructions. Quantitative PCR was performed using the TaqMan Gene Expression Master Mix (Thermofisher, Cat.#4369016) corresponding to TaqMan Gene Expression Assay Probes (Thermofisher, Cat. # 4331182 for *Il1b,* Cat.# 4331182 for *Hprt*), and 100 ng of cDNA template per reaction in a 96-well plate. Thermal Cycling conditions were: 2 minutes at 50°C, 10 minutes at 95°C, followed by 60 cycles of 15 seconds at 95°C and 60 seconds at 60 °C. The quantitative PCRs were performed using the Bio-Rad CFX96 Real-Time PCR Detection System (RRID: SCR_018064). HPRT was used as a housekeeping gene (Ho & Patrizi, 2021).

### Nanostring analysis

#### Over-Representation Analysis

NanoString data from the brains of HFD-C-fed *Stat4^ΔLysM^Ldlr^−/−^* and *Stat4^fl/fl^Ldlr^−/−^* mice were acquired from the Gene Expression Omnibus database (accession number GSE240045, RRID:SCR_005012) (Zhang et al., 2023). 24-week-old mice were sustained on the HFD-C diet for 16 weeks before sacrifice and tissue RNA extraction. Significantly upregulated or downregulated (p<0.05) (log2Fold>|1.5|) genes underwent pathway analysis. Kyoto Encyclopedia of Genes and Genomes (KEGG) Pathway analysis was performed using SRPlot (RRID:SCR_025904). Significantly enriched pathways (p_adj_.<0.05) were selected and graphed using a Sankey and dot plot in SRplot.(Tang et al. 2023)

#### Gene Set Enrichment Analysis

NanoString data (accession number GSE240045, RRID:SCR_005012) (Zhang et al., 2023) was additionally interrogated using Gene Set Enrichment analysis in Webgestalt (Elizarraras et al., 2024) using Reactome Pathway analysis. Pathways represented in the Enrichment dot plot (produced in SRplot), were selected from those that were statistically significantly enriched (p<0.05).(Tang et al. 2023)

### Efferocytosis Assay

The thymus was dissected, and a thymocyte cell suspension was obtained using a 70 μm mesh. Apoptosis was then induced through UV irradiation as described in Shi et al. (Shi et al., 2022). Cells were then stained with CellTrace Far Red (Thermofisher, Cat.#C34564) per manufacturer’s instructions. Bone marrow was isolated from the bones of C57BL/6 mice, filtered using a 70 μm mesh, and red blood cells were lysed using ACK Lysis buffer (150mM ammonium chloride, 10mM potassium bicarbonate, 0.1 mM EDTA). Cells were resuspended in HBSS, and neutrophils were isolated using the Stem Cell EasySep Mouse Neutrophil Enrichment Kit (Stem Cell, Cat# 19762) following the manufacturer’s instructions. Neutrophils were plated in RPMI with 10% FBS overnight at 37°C. The next day, the cells were washed with HBSS and then stained with CFSE (Thermofisher, Cat# C34554) at 1:40,000 for 20 minutes at 37°C. Neutrophils were washed with HBSS with 10% HBSS. 0.5×10^6^ cells from a single cell suspension of mouse brains were plated in a 6-well plate in 3 mL of RPMI with 10% FBS. 0,1×10^6^ stained, aged neutrophils or apoptotic thymocytes were added to each well, and the plate was incubated at 37°C for 1 hour. The cell suspension was collected and stained with Abs for FACS analysis.

### Immunohistochemistry

Brains were formalin-fixed and paraffin-embedded before being sagittal sectioned into 5 µm slices. Slides were heated at 58°C overnight prior to deparaffinization and rehydration in xylenes and ethanol of decreasing concentration. We performed antigen unmasking using a citrate-based antigen unmasking solution (Vector Labs H-3300-250, RRID: AB_2336226) in a microwave for 20 minutes, followed by 20 minutes at room temperature. Samples were incubated in peroxidase block (Vector Labs, Cat.# SP-6000-100, RRID:AB_2336257) for 30 minutes, followed by 5% BSA in tris-buffered saline with 0.05% Tween 20 (TBST) for 30 minutes. Slides were incubated with an antibody targeting pTau (Ser199) (Invitrogen Cat.# 44734G, RRID: AB_2533737) at 1:1000 for 1 hour before incubating with goat anti-rabbit biotin (1:200) (Vector Labs, Cat# BA-1000, RRID: AB_2313606) for 30 minutes. Slides were incubated with ABC (Vector Labs, Cat# PK-7100, RRID: AB_2336827) for 30 minutes and then with DAB (Vector Labs, Cat# SK-4105, RRID: AB_2336520) for 10 minutes. They were then dipped in water, counterstained with hematoxylin (Vector, Cat# H-3401, RRID: AB_2336842), and subsequently dehydrated before being mounted. Slides were imaged at 40x magnification on a Nikon Coolscope, and the percentage of positive area across the sagittal section 10 brain regions was averaged using ImageJ (RRID:SCR_003070). Representative images were taken on a Zeiss Observer Z7, white balanced, and brightness/contrast were adjusted in ZenBlue.

### Statistical Analysis

Data are represented as mean±SD. Outliers were identified using the mean ± 1.5x SD and excluded if these criteria were not met. For the efferocytosis assay, total CFSE+ cell numbers were normalized by dividing each number by the average for the *Stat4^fl/fl^Ldlr^−/−^* HFD-C mice of the cohort. Statistical analysis was performed using Prism 10 (Graphpad, RRID:SCR_002798). Sample size was determined using a power calculation with α=0.05 and β=0.2. Chow-fed animals were limited in number. Males and females were analyzed separately. Normality of the data was determined using the Shapiro-Wilk test, and the Forsythe test was employed to assess equality of variances. Comparisons between groups were performed by one-way ANOVA with a Šídák multiple comparisons test for normally distributed data with equal variances, by Brown-Forsythe and Welch ANOVA with a Dunnett’s 3T correction for normally distributed data with unequal variances, or by Kruskal-Wallis with Dunn’s correction for data that is not normally distributed. For the novel object recognition test, a one-sample t-test compared to 50 was performed to determine whether the recognition index was greater than chance.

## 3. Results

### STAT4-deficiency reduces cognitive decline in HFD-C-fed atherosclerotic *Ldlr^−/−^* mice

Epidemiological studies link vascular disease risk factors, including atherosclerosis, obesity, and hypertension, with neurodegeneration. IL-12 modulates neuroinflammation and plays a proatherogenic role by activating STAT4. To determine the role of LysM^Cre^-derived STAT4 in regulating memory, we performed several behavioral tests following 28 weeks of HFD-C diet feeding (Fig.1A). HFD-C diet affects cholesterol and aortic root atherosclerosis in male and female *Ldlr^−/−^* mice differently (Mansukhani et al., 2017). Therefore, male and female mice were evaluated separately to determine whether the effects of *Stat4* knockout differed between sexes.

**Figure 1.**
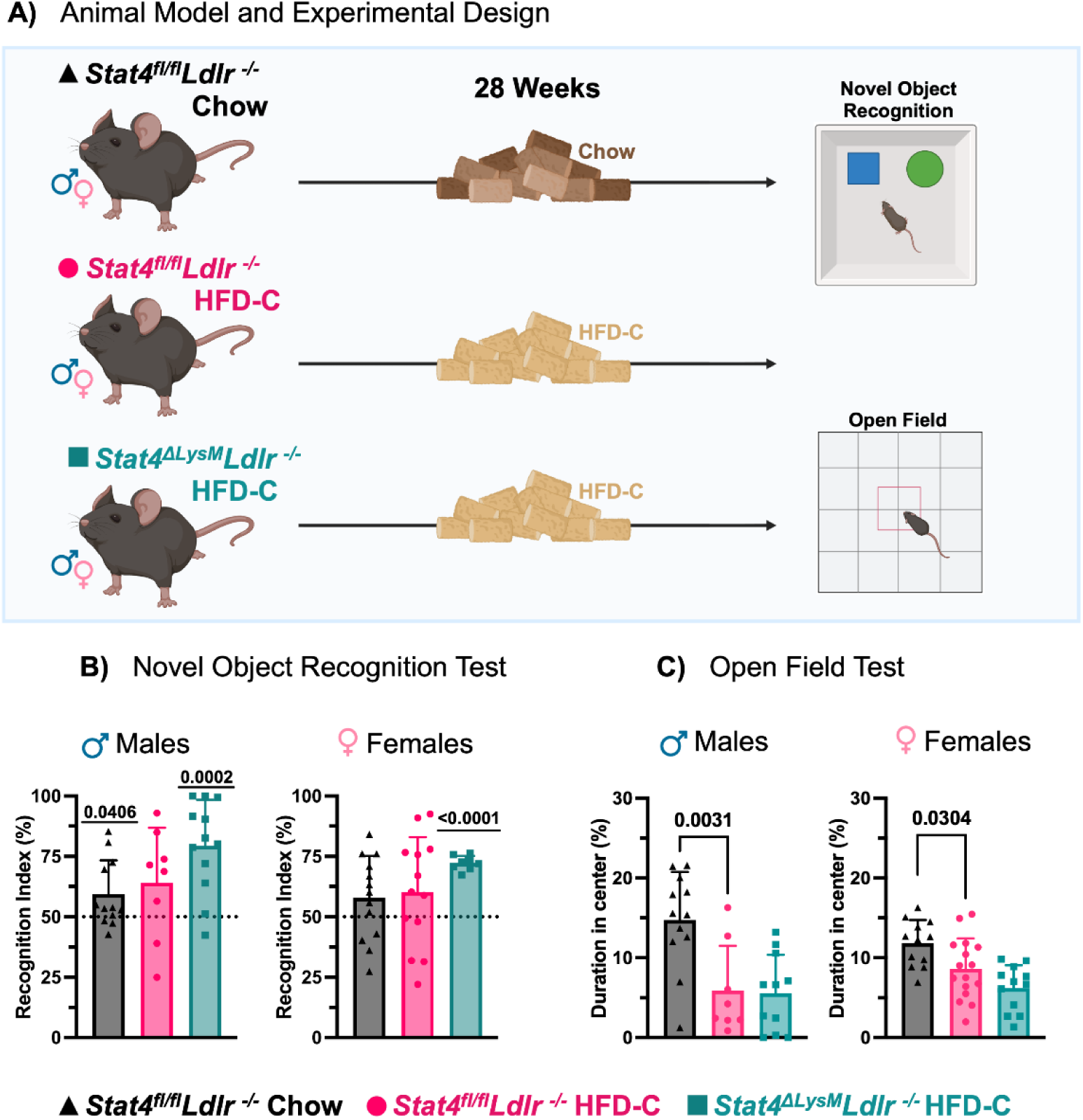
Myeloid-derived Stat4 drives cognitive decline in Ldlr^−/−^ mice. **(A)** 8-12-week-old *Stat4^fl/fl^Ldlr^−/−^* and *Stat4^ΔLysM^Ldlr^−/−^* mice were kept on either HFD-C or regular chow diet for 28 weeks before undergoing novel object recognition and open field behavioral testing. **(B)** A recognition index was calculated for the novel-object recognition test. Both male and female *Stat4^ΔLysM^Ldlr^−/−^* mice demonstrate a significant preference for the novel object after 28 weeks of HFD-C. Male chow-fed *Stat4^fl/fl^Ldlr^−/−^* mice also show a significant preference for the novel object, but female mice do not. One-sample t-test (compared to 50%). **(C)** The percentage of time spent in the center is calculated using open-field test data. Male HFD-C-fed *Stat4^ΔLysM^Ldlr^−/−^* mice spend less time in the center of the open field than 28-week and 28-week chow-fed *Stat4^fl/fl^Ldlr^−/−^* mice. Female 28-week HFD-C-fed *Stat4^ΔLysM^Ldlr^−/−^* mice spend reduced time in the center vs chow-fed *Stat4^fl/fl^Ldlr^−/−^* females. One-way ANOVA with Šídák multiple comparisons (n = 8-12 for males and 9-16 for females/per group).

NOR was used to assess long-term memory (Lueptow, 2017). Recognition index was calculated as the percentage of time spent exploring the novel object relative to the total exploration time of both the familiar and novel objects. This index reflects the animal’s ability to discriminate and recognize the novel object, with higher values indicating better memory or recognition performance. A recognition index of 50% represents a chance level, meaning the animal spends equal time exploring both objects and shows no preference. Animals that spent significantly more time with the novel object than the familiar one on day two of testing (resulting in a recognition index above 50%) demonstrate better long-term memory than those performing at chance level (Lueptow, 2017).

For age-matched female mice fed a chow diet, there was no difference in interaction compared to chance (Fig.1B). Age-matched male *Stat4^fl/fl^Ldlr^−/−^* mice fed a standard chow diet had a significantly higher interaction with the novel object compared to chance. Both male and female *Stat4^fl/fl^Ldlr^−/−^* mice fed HFD-C diet for 28 weeks had memory impairments as demonstrated by a recognition index that was not significantly different from chance. However, both male and female *Stat4^ΔLysM^Ldlr^−/−^* mice fed an HFD-C diet for 28 weeks had a significantly higher interaction with the novel object compared to chance (Fig.1B). Together, these results indicate that impairments in long-term memory were ameliorated by LysM^Cre^-specific deletion of *Stat4*.

An additional feature of cognitive impairments is anxiety. To evaluate whether LysM-specific deletion of STAT4 affects anxiety, we performed Open Field tests after 28 weeks on an HFD-C diet. Animals that spent more time in the center portion of a 16in x 16in container compared to the duration spent on the periphery of the container were considered less anxious than those who spent more time in the periphery. In line with previous reports (Freeman et al., 2014), HFD-C feeding clearly induced more anxiety in *Ldlr^−/−^* males and females after prolonged HFD-C diet, as determined by reduced duration spent in the center (Fig.1C). Interestingly, STAT4 deficiency did not attenuate anxiety levels in HFD-C-fed *Stat4^ΔLysM^Ldlr^-−/−^* vs HFD-C STAT4-sufficient *Ldlr^−/−^* mice, demonstrating the dominant effect of HFD-C feeding on the regulation of anxiety levels.

### LysM^Cre^-specific deletion of *Stat4* ameliorates phospho-tau expression in the brain

Tau hyperphosphorylation is a significant component of the pathological process in various tauopathies, leading to toxic tau aggregation, disruption of cellular functions, and promotion of neurodegeneration (Neddens et al., 2018). Several studies demonstrate that an HFD regimen induces Tau hyperphosphorylation via GSK3β activation in C57BL/6 mice (Bhat & Thirumangalakudi, 2013; Liang et al., 2023). Increased phosphorylation of Tau at the Ser199 site has been reported in other models of dyslipidemia (Lénárt et al., 2012). To further understand the mechanism behind behavioral abnormalities associated with HFC-C intake, we examined the expression of pTau in sagittal sections of the brains of STAT4-sufficient and STAT4-deficient *Ldlr^−/−^* mice (Fig.2A). While pTau was not statistically increased after 28-week HFD-C feeding in the brains of *Stat4^fl/fl^Ldlr^−/−^* mice, HFD-C-fed *Stat4^fl/fl^Ldlr^−/−^* males demonstrated a moderate increase in pTau in comparison with chow-fed *Stat4^fl/fl^Ldlr^−/−^* males (Fig.2B). Importantly, the LysM^Cre^-specific deletion of *Stat4* decreased the percent positive area for pTau in the brain of both male and female HFD-C-fed *Stat4^ΔLysM^Ldlr^−/−^* mice in comparison with HFD-C-fed *Stat4^fl/fl^Ldlr^−/−^* animals, suggesting a role for LysM^Cre^-derived STAT4 in Tau hyperphosphorylation (Fig.2).

**Figure 2.**
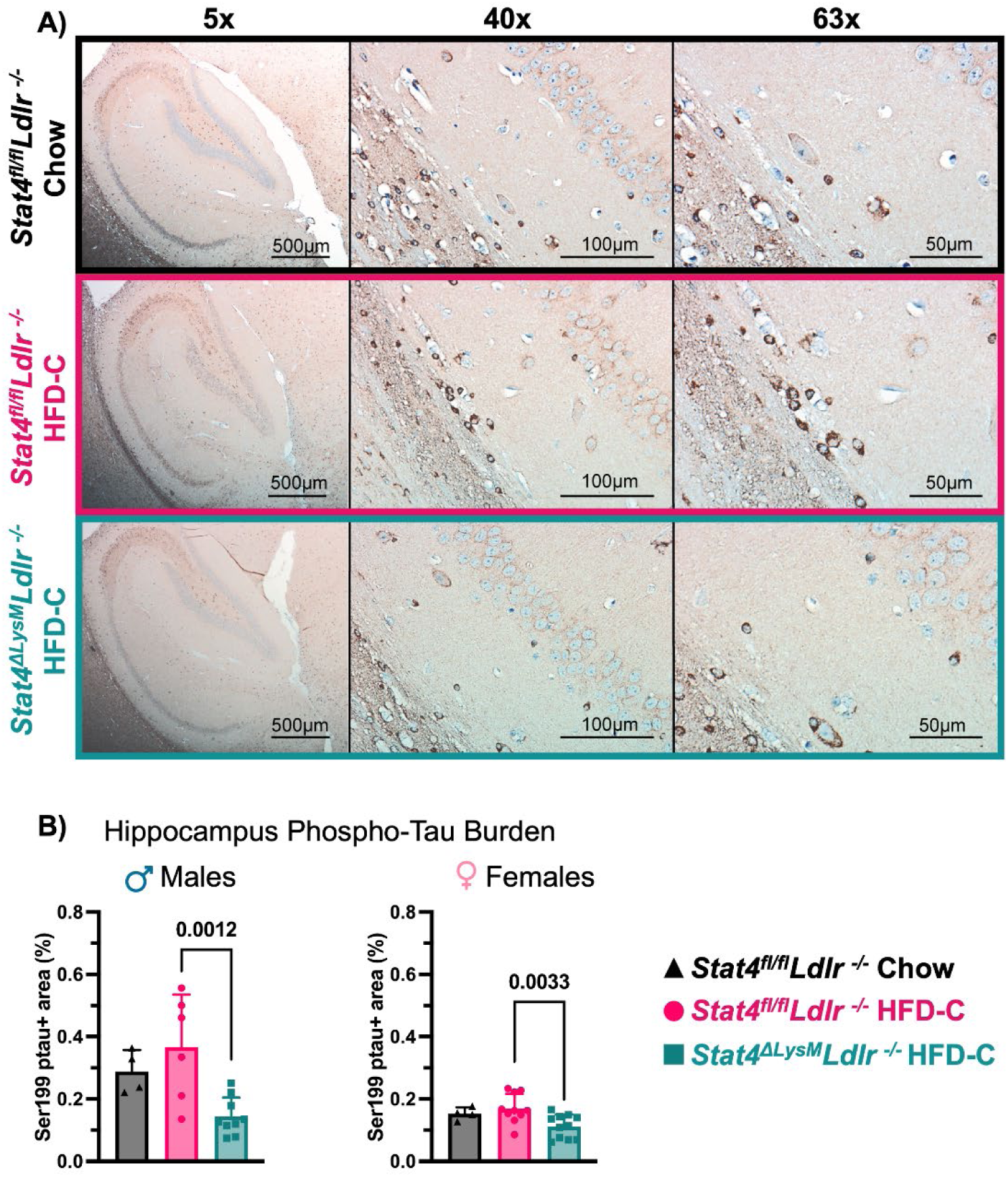
LysM^Cre^-specific deletion of Stat4 ameliorates pTau expression in the brain. 8-12-week-old *Stat4^fl/fl^Ldlr^−/−^* and *Stat4^ΔLysM^Ldlr^−/−^* mice were kept on HFD-C or chow diet for 28 weeks. **(A)** Representative imaging of Ser199-phosphorylated tau in the hippocampus at 5x, 40X, and 63x. **(B**) Percent of positive area for Ser199-phosphorylated Tau in the brain of male and female HFD-C-fed or chow diet-fed *Stat4^ΔLysM^Ldlr^−/−^* and *Stat4^fl/fl^Ldlr^−/−^* mice. One-way ANOVA with Šídák multiple comparisons, (n=4-10/per group).

### LysM^Cre^-specific deletion of *Stat4* attenuated microglia numbers and activation in HFD-C-fed *Ldlr^−/−^* mice

Having demonstrated that LysM^Cre^-driven STAT4 deficiency improves cognitive function in *Stat4^ΔLysM^Ldlr^−/−^* mice, we sought to corroborate our findings by evaluating neuroimmune-mediated mechanisms using spectral flow cytometry. Long-term exposure to HFD has previously been associated with hypothalamic microgliosis, characterized by increased numbers and activation of CD45^low^CD11b^+^ microglia and CD45^hi^CD11b^+^ monocyte infiltration (Valdearcos et al., 2017). In line with previous reports, HFD-C feeding induced multiple effects on the immune cell populations in the brains. HFD-C feeding moderately increased the number of CD45^+^ cells in the brains of HFD-C-fed *Stat4^fl/fl^Ldlr^−/−^* mice (Fig.3A). Importantly, LysM^Cre^-specific deletion of *Stat4* significantly reduced the number of CD45^+^ cells in the brains of HFD-C-fed *Stat4^ΔLysM^Ldlr^−/−^* vs HFD-fed *Stat4^fl/fl^Ldlr^−/−^* mice (Fig.3A).

**Figure 3.**
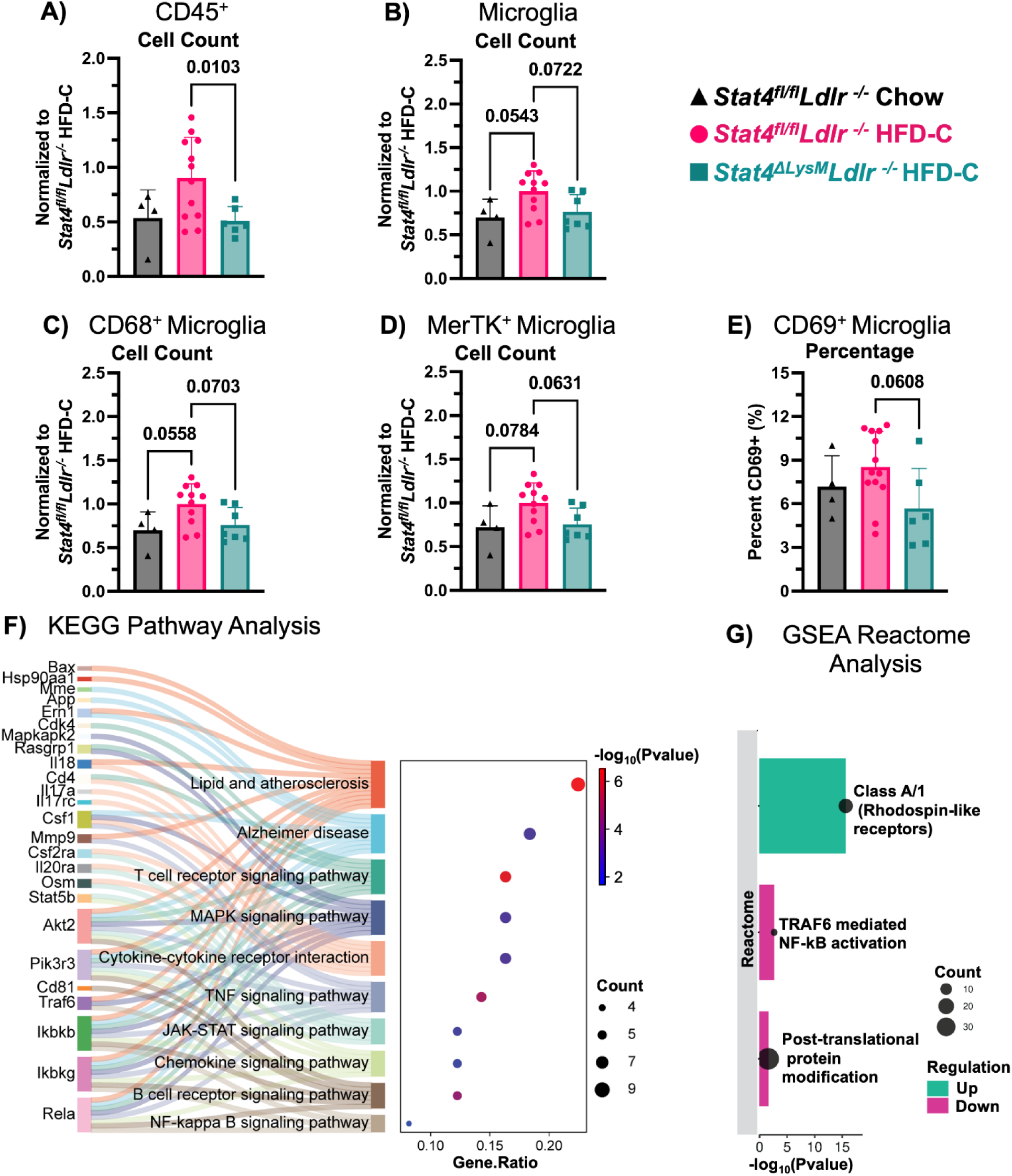
LysM^Cre^-specific deletion of Stat4 attenuated microglia numbers and activation in HFD-C-fed Ldlr^−/−^ mice. 8-12-week-old *Stat4^fl/fl^Ldlr^−/−^* and *Stat4^ΔLysM^Ldlr^−/−^* mice were kept on either HFD-C or chow diet for 28 weeks. At sacrifice, brains were mechanically digested and stained for flow cytometry. **(A-D)** Relative ratios of total cell numbers are calculated to HFD-C-fed *Stat4^fl/fl^Ldlr^−/−^* cell numbers. Each data point is normalized by dividing the average value of HFD-C-fed *Stat4^fl/fl^Ldlr^−/−^* mice collected the same day. **(A)** Ratios of CD45^+^ cells in the brains of the indicated groups. HFD-C-fed *Stat4^ΔLysM^Ldlr^−/−^* mice have significantly fewer CD45^+^ cells in the brain compared to HFD-C-fed *Stat4^fl/fl^Ldlr^−/−^* mice. **(B)** Ratios of CD45^low^Tmem119^+^Ly6G^-^ microglia are shown. HFD-C feeding increases microglia numbers in HFD-C-fed *Stat4^fl/fl^Ldlr^−/−^* mice compared to chow-fed *Stat4^fl/fl^Ldlr^−/−^* mice, whereas LysM^Cre^-specific deletion of *Stat4* reduces this effect. **(C)** Ratios of CD45^low^Tmem119^+^Ly6G^-^ microglia expressing CD68 are shown. HFD-C-fed *Stat4^fl/fl^Ldlr^−/−^* mice have more CD68^+^ microglia compared to both HFD-C-fed *Stat4^ΔLysM^Ldlr^−/−^* mice and chow-fed *Stat4^fl/fl^Ldlr^−/−^* mice. **(D)** Ratios for MerTK expression by microglia. HFD-C-fed *Stat4^ΔLysM^Ldlr^−/−^* mice have fewer MerTK+ microglia compared to HFD-C-fed *Stat4^fl/fl^Ldlr^−/−^* mice. **(E)** CD69 expression by microglia. HFD-C-fed *Stat4^ΔLysM^Ldlr^−/−^* mice display a decreased percentage of CD69^+^ microglia compared to HFD-C-fed *Stat4^fl/fl^Ldlr^−/−^* mice. One-way ANOVA with Šídák multiple comparisons or Brown-Forsythe and Welch ANOVA with Dunnett’s T3 multiple comparisons (n=4-12/per group). **(F)** A Sankey plot shows KEGG pathways down-regulated in HFD-C-fed *Stat4^ΔLys^MLdlr^−/−^* mice compared to *Stat4^fl/fl^Ldlr^−/−^* mice (FDR < 0.05), identified using SRPlot. The dot size is based on the gene count enriched in the pathway. **(G)** Enrichment Plot shows Reactome Pathway analysis using GSEA, identified using WebGestalt and graphed using SRplot. The dot size indicates the number of genes associated with the indicated Reactome pathway.

The number of CD45^low^Tmem119^+^Ly6G^-^ microglia was also elevated in HFD-C-fed vs. chow diet *Stat4^fl/fl^Ldlr^−/−^* mice (Fig.3B), suggesting active microgliosis, a state of activation and increased proliferation of microglia, that is often seen in AD and other forms of cognitive decline. STAT4 deficiency attenuated the detrimental effects of atherogenesis and HFD-C feeding, as we detected a reduced number of microglia in HFD-C-fed *Stat4^ΔLysM^Ldlr^−/−^* vs *Stat4^fl/fl^Ldlr^−/−^* mice (Fig.3B, p<0,07).

CD68 is an established marker for activated microglia in the context of neuroinflammation and cognitive decline (Farso et al., 2013). In correlation with the increased microglia numbers, the relative number of microglia expressing CD68^+^ was significantly altered between the groups, with a trend toward increased CD68^+^ microglia in *Stat4^fl/fl^Ldlr^−/−^* mice fed HFD-C compared to chow, and a decrease in HFD-C-fed *Stat4^ΔLysM^Ldlr^−/−^* vs *Stat4^fl/fl^Ldlr^−/−^* mice (Fig.3C). MerTK plays a critical role in the clearance of apoptotic cells during efferocytosis, a process closely tied to microglial activation state in AD. In line with the elevated number of CD68^+^ microglia, the number of MerTK+ microglia was increased under pressure of HFD-C and atherogenesis in HFD-C-fed *Stat4^fl/fl^Ldlr^−/−^* mice (Fig.3D, p<0.06). Notably, the number of MerTK^+^ microglia was reduced in HFD-C-fed *Stat4^ΔLysM^Ldlr^−/−^* mice in comparison with age and diet-matched *Stat4^fl/fl^Ldlr^−/−^* controls (Fig.3D). Although CD69 is not a typical marker expressed by microglia, hypercholesterolemia and associated atherosclerosis increased the percentage of CD69-expressing CD45^low^CD11b^+^ microglia in HFD-C *Stat4^fl/fl^Ldlr^−/−^* mice. This increase was significantly reduced in HFD-C-fed *Stat4^ΔLysM^Ldlr^−/−^* mice (Fig.3E). Thus, LysM^Cre^-driven STAT4 deficiency protects against HFD-induced atherosclerosis-associated neuroinflammation by reducing microgliosis and numbers of activated CD68^+^, MerTK^+^, and CD69^+^microglia, thereby dampening detrimental hyperactivation of microglia.

Previous studies have performed NanoString analysis on the brains of *Stat4^ΔLysM^Ldlr^−/−^* vs *Stat4^fl/fl^Ldlr^−/−^* mice fed an HFD-C for 16 weeks (Zhang et al., 2023). Using these publicly available DEGs, we performed a KEGG pathway analysis to identify relevant pathways downregulated in the brains of *Stat4^ΔLysM^Ldlr^−/−^* mice compared to those of *Stat4^fl/fl^Ldlr^−/−^* mice during HFD-C-induced atherosclerosis. Predictably, several genes involved in the JAK/STAT signaling pathway were decreased. Additionally, several genes related to atherosclerosis, AD, TNFα signaling, NF-κB signaling, and chemokine signaling were downregulated in the *Stat4^ΔLysM^Ldlr^−/−^* mice compared to *Stat4^fl/fl^Ldlr^−/−^* mice (Fig.3F). In summary, our data implicates LysM^Cre^-driven STAT4 expression as a critical regulator of inflammation in the brain in conditions of atherosclerosis.

We sought to further investigate the effects of STAT4 deficiency in this NanoString data using gene set enrichment analysis (GSEA) to capture any subtle, but coordinated, changes in pathways that may not be apparent in the over-representation analysis performed in Fig.3F. Reactome pathway analysis using GSEA further supported that NF-κB signaling was attenuated in the brains of *Stat4^ΔLysM^Ldlr^−/−^* vs *Stat4^fl/fl^Ldlr^−/−^* mice during HFD-C (Fig.3G). Additionally, the brains of *Stat4^ΔLysM^Ldlr^−/−^* mice had increased signaling in Class A/1 (Rhodopsin-like receptors compared to *Stat4^fl/fl^Ldlr^−/−^* mice (Fig.3G). Rhodopsin-like receptors play an integral role in synaptic plasticity and memory (Wong et al., 2023). Among the genes upregulated with STAT4 deficiency in this pathway, *Ccl1* and *Cxcl16* play critical roles in myeloid chemotaxis and neuronal synaptic transmission. Notably, CCL1 is decreased in the brains of APP/PS1 AD mice.(Jorda et al., 2019) CCL1 is expressed in neurons and other brain cells and promotes chemotaxis of mononuclear cells (Jorda et al., 2019; Trebst et al., 2003). CXCL16 enhances spontaneous release and reduces action potential-dependent release of GABA at inhibitory synapses in the hippocampus, and enhances glutamate release at excitatory synapses (Di Castro et al., 2016). Interestingly, CXCL16 appears to have a neuroprotective effect, preventing excitotoxic cell death in hippocampal neurons (Rosito et al., 2012). This analysis reveals that STAT4 deficiency affects Rhodopsin-like receptors involved in immune remodeling and drives neuronal protection against excitotoxic cell death.

Activation of the post-translational protein modification signaling pathway also decreased in STAT4-deficient mice. Specifically, STAT4 deficiency decreased expression of multiple ubiquitination-related genes, including *Cul1, Uba52, Ube2n, Psmb8,* and *Psmb10.* This indicates that *LysM-* mediated STAT4 plays a role in increasing ubiquitination, which has previously tied to the accumulation of pathological neurofibrillary tangles.(Li et al., 2021; Yang et al., 2025)

### Effects of atherosclerosis and hypercholesterolemia on the brain’s immune composition

Monocytes and monocyte-derived macrophages play a complex role in neuroinflammation-driven pathologies, as these cells actively eliminate Aβ, pTau, and damaged cells, while simultaneously triggering increased inflammation through the release of proinflammatory cytokines such as IL-1α/β, IL-6, TNFα, IL-8, and TGFβ (Feng et al., 2011). Thus, emigrated monocytes play a dual role in the brain, either exacerbating or alleviating disease progression. Our results demonstrate that HFD-C feeding and atherosclerotic conditions altered the composition of myeloid cell subsets in the brain of HFD-C vs Chow diet-fed *Stat4^fl/fl^Ldlr^−/−^* mice, with a significant decrease in the number of CD68^+^MHC-II^+^CD206^+^ myeloid cells (Fig.4A), which represent phagocytic, antigen-presenting myeloid cells with anti-inflammatory potential.

**Figure 4.**
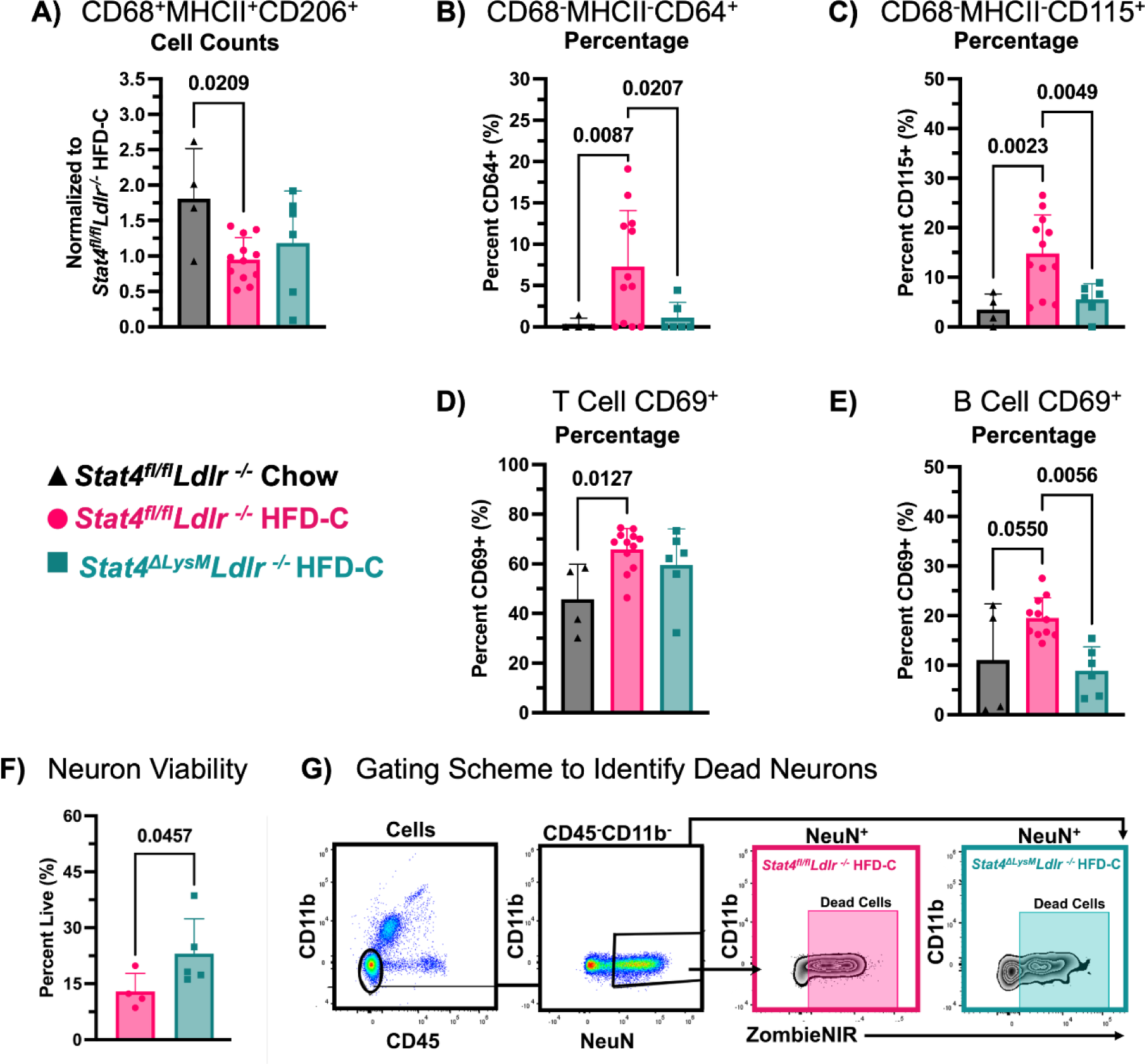
Effects of atherosclerosis and hypercholesterolemia on the immune composition of the brain. 8-12-week-old *Stat4^fl/fl^Ldlr^−/−^* and *Stat4^ΔLysM^Ldlr^−/−^* mice were kept on either HFD-C or chow diet for 28 weeks. At sacrifice, brains were mechanically digested and stained for flow cytometry. **(A)** Relative ratios of total cell numbers to HFD-C-fed *Stat4^fl/fl^Ldlr^−/−^* cell numbers are calculated. Each data point is normalized by dividing the average value of HFD-C-fed *Stat4^fl/fl^Ldlr^−/−^* mice collected the same day. HFD-C-fed *Stat4^fl/fl^Ldlr^−/−^* mice have significantly decreased CD206^+^CD68^+^MHCII^+^ cells compared to chow-fed *Stat4^fl/fl^Ldlr^−/−^* mice. **(B, C**) Percentages of CD64^+^ and CD115^+^ expressing CD68^-^MHCII^-^ cells are increased with HFD-C and are ameliorated in *Stat4^ΔLysM^Ldlr^−/−^* mice. **(D)** HFD-C increases the percentage of CD3^+^ CD69^+^ T cells in *Stat4^fl/fl^Ldlr^−/−^* mice. **(E)** HFD-C-feeding reduces the percentage of CD69^+^ B cells in the brain of *Stat4^fl/fl^Ldlr^−/−^* mice (n=4-12/per group). **(F)** Percent of Live (ZombieNir^-^) neurons in brains of HFD-C-fed *Stat4^fl/fl^Ldlr^−/−^* and *Stat4^ΔLysM^Ldlr^−/−^* mice. **(G)** Representative gating strategy for neuronal viability. Brains were initially gated on single cells to exclude doublets and then gated on cell populations to exclude debris (n=4-5/per group). One-way ANOVA with Šídák multiple comparisons or Brown-Forsythe and Welch ANOVA with Dunnett’s T3 multiple comparisons.

HFD-C feeding significantly increased the percentage of CD64-expressing CD68^-^MHC-II^-^cells, whereas LysM^Cre^-driven STAT4 deficiency, in turn, decreased this percentage of immature CD68-MHC-II-CD64^+^ monocytes in the brain of *Stat4^ΔLysM^Ldlr^−/−^* mice (Fig.4B). These results suggest that HFD-C feeding increases immature monocyte levels in a STAT4-dependent manner. CD115 is the receptor for macrophage colony-stimulating factor (M-CSF), essential for monocyte survival, proliferation, and differentiation. CD115^+^ immature monocytes are critical during the early phases of inflammation and tissue infiltration, before full activation and maturation into macrophages or antigen-presenting cells.(Monaghan et al., 2019) CD115 expression was increased by HFD-C feeding, further suggesting that atherogenesis supports monocyte maturation in the brain, and that this effect is ameliorated by STAT4 deficiency (Fig.4C).

Beyond myeloid cells, lymphoid cells also regulate neuroinflammation. Mice with tauopathy display an increased number of T cells that correlates with the extent of neuronal loss (Chen et al., 2023). As our mouse model of atherosclerosis and HFD-induced obesity exhibits some signatures of cognitive decline, we next examined the presence of T cells in the brain. Brains of HFD-C- vs chow-diet-fed *Stat4^fl/fl^Ldlr^−/−^* mice displayed an increased expression of CD69 on CD3^+^ cells (Fig.4D), indicating a recent activation of this population. These alterations in the number and phenotype of T cells were surprisingly STAT4-independent. The role of B cells in regulating neuroinflammation remains under investigation; however, several reports suggest an immunoglobulin isotype-specific role. Interestingly, brains from HFD-C-fed *Stat4^fl/fl^Ldlr^−/−^* mice had a moderate increase in the percentage of CD69^+^CD19^+^ B cells, and a significant reduction of activated CD69^+^ B cells was observed in *Stat4^ΔLysM^Ldlr^−/−^* mice (Fig.4E). To explore the impact of these immune composition and activation changes on neuronal health, we investigated the viability of NeuN^+^ cells in the brains of HFD-C fed *Stat4^fl/fl^Ldlr^−/−^* and *Stat4^ΔLysM^Ldlr^−/−^* mice. Neurons from the brains of *Stat4^ΔLysM^Ldlr^−/−^* mice had a higher viability percentage compared to *Stat4^fl/fl^Ldlr^−/−^* mice, indicating that STAT4 deficiency improved local microenvironments that support neuronal health (Fig.4F,G). Altogether, our data demonstrate substantial changes in brain immune composition, activation, and functions induced by HFD-C feeding and atherosclerosis, and partially regulated by LysM^Cre^-dependent STAT4 expression.

### LysM^Cre^-specific deletion of *Stat4* **i**ncreased microglial efferocytosis

Macrophages maintain systemic self-tolerance and promote the resolution of inflammation and tissue repair through efferocytosis. It has been previously demonstrated that *Ldlr^−/−^* mice express increased markers of apoptosis in the brain, colocalized with neuronal markers, suggesting that cell death may play a role in the neurological deficits in *Ldlr^−/−^* mice (de Oliveira et al., 2020). Since macrophages and microglia play a crucial role in efferocytosis, we investigated whether LysM^Cre^-specific *Stat4* knockout affects the clearance of apoptotic cells. Apoptotic neutrophils or thymocytes were used to evaluate efferocytosis in immune cells isolated from the brains of these animals by FACS analysis (Fig.5A). HFD-C moderately increased the total number of microglia efferocytosis in *Stat4^fl/fl^Ldlr^−/−^* mice (Fig.5B). Although HFD-C feeding resulted in moderately increased numbers of microglia, the percentage of microglia undergoing efferocytosis (CFSE^+^) was further increased in *Stat4^fl/fl^Ldlr^−/−^* mice (Fig.5D). Notably, the relative proportion of microglia that was actively involved in efferocytosis was further amplified in *Stat4^ΔLysM^Ldlr^−/−^* mice (Fig.5B and 5D). The percentage of efferocytotic monocytes and macrophages, identified as CD45^+^Tmem119^-^CD11b^+^CFSE^+^ cells, was significantly increased with HFD-C feeding, but not further affected by *Stat4* deficiency (Fig.5E). However, the overall total number of CD11b^+^ cells that were positive for efferocytosis was significantly increased in *Stat4^ΔLysM^Ldlr^−/−^* mice in comparison with *Stat4^fl/fl^Ldlr^−/−^* mice (Fig.5C). Thus, these results suggest that STAT4 plays a crucial role in regulating the efferocytotic activity of microglia. PCR of brain lysates also revealed decreased *Il1b* expression in the brains of mice with LysM^Cre^-specific *Stat4* knockout, suggesting reduced pro-inflammatory signaling (Fig.5F).

**Figure 5.**
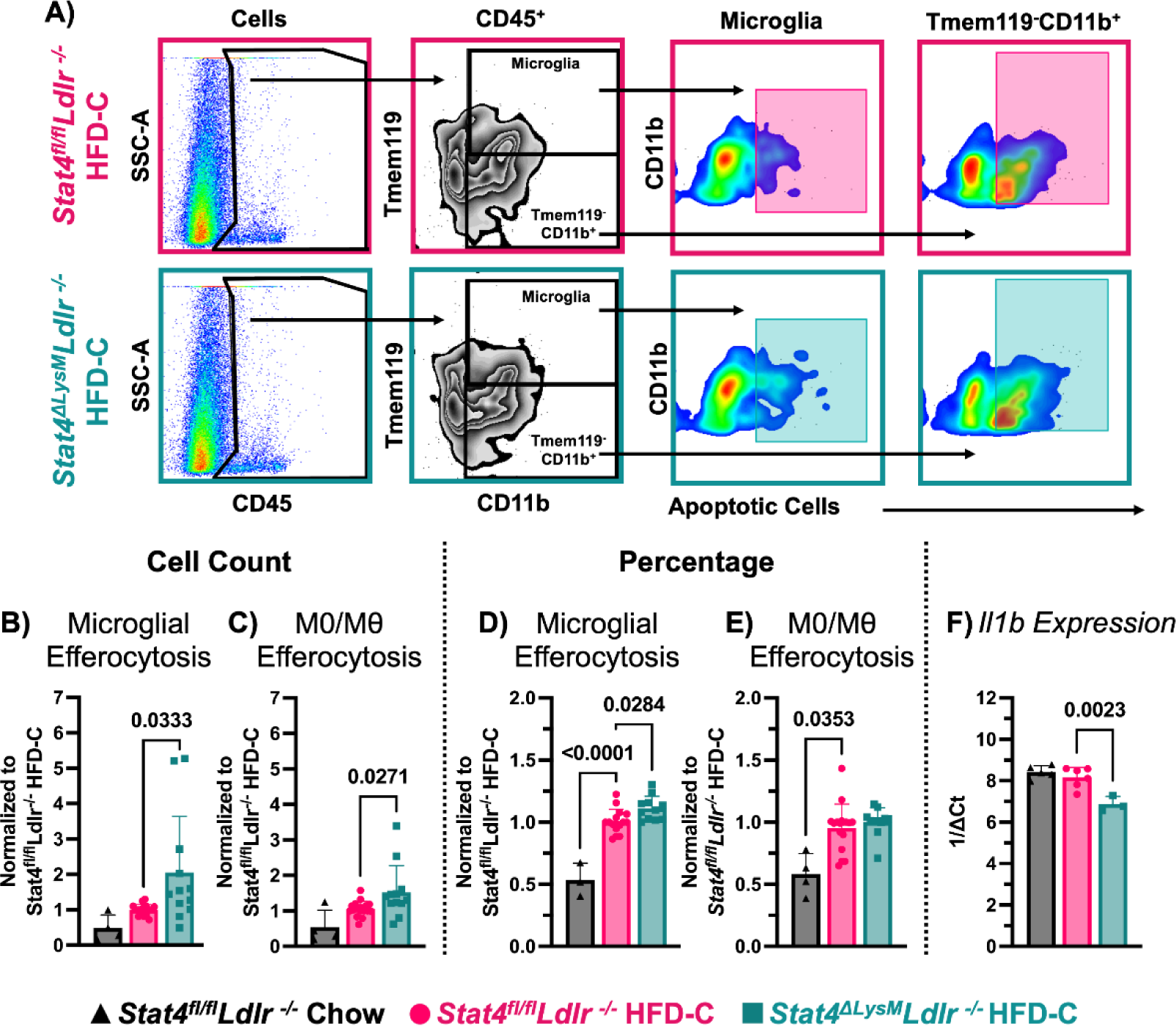
LysM^Cre^-specific deletion of Stat4 increases microglial efferocytosis. 8-12-week-old *Stat4^fl/fl^Ldlr^−/−^* and *Stat4^ΔLysM^Ldlr^−/−^* mice were kept on HFD-C for 16-28 weeks. 0.5×10^6^ cells were isolated from the brain by mechanical digestion and incubated with 0,1×10^6^ aged, CFSE-stained aged neutrophils or CellTrace Far Red-stained apoptotic thymocytes for 1 hour. The cells were then stained and analyzed by flow cytometry. **(A)** Representative gating strategy for efferocytosis assay. **(B-E)** Relative ratios of total efferocytosis+ microglia and efferocytosis+ CD45^+^CD11b^+^ myeloid cell numbers (B-C) and percentage (D-E) to HFD-C-fed *Stat4^fl/fl^Ldlr^−/−^* cell numbers. Each data point is normalized by dividing the average value for HFD-C-fed *Stat4^fl/fl^Ldlr^−/−^* mice collected the same day. (**B-C**) Ratios of total cell numbers of efferocytosis^+^ cells are shown. HFD-C-fed *Stat4^ΔLysM^Ldlr^−/−^* mice have significantly more efferocytotic CD11b^+^ myeloid cells in the brain than HFD-C-fed *Stat4^fl/fl^Ldlr^−/−^* mice or HFD-C-fed *Stat4^ΔLysM^Ldlr^−/−^* mice. **(D-E)** Ratios of percentages of efferocytosis+ cells out of the total CD45^+^ cells are shown. (D) HFD-C increases the percentage of efferocytotic microglia in the brains of *Stat4^fl/fl^Ldlr^−/−^* mice, while HFD-C-fed *Stat4^ΔLysM^Ldlr^−/−^* mice have a further increased percentage of CFSE^+^ microglia in the brain compared to *Stat4^fl/fl^Ldlr^−/−^* mice fed a HFD-C. (E) HFD-C feeding increases the percentage of efferocytotic Tmem119^-^CD11b^+^ cells in the brains of *Stat4^fl/fl^Ldlr^−/−^* mice, (n=4-12/per group) (F) *Il1b* gene expression in the brains of HFD-C *Stat4^ΔLysM^Ldlr^−/−^* mice vs HFD-C-fed *Stat4^fl/fl^Ldlr^−/−^* mice, (n=4-6/per group). One-way ANOVA with Šídák multiple comparisons or Kruskal-Wallis with Dunn’s multiple comparisons.

## 4. Discussion

Neurodegenerative diseases and atherosclerosis are characterized by common signatures of vascular dysfunction and chronic inflammation, and are accelerated with aging (Stahr & Galkina, 2022; Xie et al., 2020). Neuroinflammation is a critical component of neurodegenerative diseases, driving chronic activation of microglia and astrocytes within the central nervous system. Pro-inflammatory cytokines, including IL-1β, IL-6, and TNFα, are elevated in the brain during neuroinflammation, promoting neuronal damage and synaptic dysfunction and exacerbating Aβ and Tau pathologies. Systemic inflammation can disrupt the blood-brain barrier, allowing leukocyte infiltration into the CNS and intensifying neuroinflammation. Surprisingly, little is known about how atherosclerosis influences neurodegeneration and cognitive health.

In this study, we show that atherosclerosis and HFD-C feeding induce cognitive decline, as evidenced by decreased performance in the NOR and Open Field tests. These data strongly support a line of epidemiological evidence connecting atherosclerosis progression with various pathologies associated with neurodegeneration. Recent data have tied cardiovascular risk factors such as ageing, hyperlipidemia, hypometabolism, hypertension, and obesity to increased risk for AD (Maciejewska et al., 2021). Additionally, the severity of atherosclerosis is associated with neuroinflammation and cognitive decline (Dolan et al., 2010; Gottesman et al., 2020; Xiang, 2017). Interestingly, our data demonstrate sexual dimorphism in responses to HFD feeding and atherosclerosis, as male *Ldlr^−/−^* mice with atherosclerosis were more affected by these conditions during the cognitive tests. These results further extend previous findings, showing that the male brain is more susceptible to metabolic changes induced by HFD-C feeding, resulting in impaired learning and memory and altered anterior cortex functions (Evans et al., 2024; Murtaj et al., 2022). Importantly, our study further reveals a potential mechanism underlying atherosclerosis-associated cognitive decline. We identify that STAT4 is a key component driving cognitive decline, as LysM^Cre^-specific STAT4 deficiency ameliorated the cognitive deficits observed in 28-week HFD-C-fed *Ldlr^−/−^* mice in the Novel Object recognition test.

STAT4 appears to mediate cognitive decline in *Ldlr^−/−^* mice through multiple, interconnected pathways. We detected a moderate decrease in pTau expression in the brains of *Stat4^ΔLysM^Ldlr^−/−^* vs *Stat4^fl/fl^Ldlr^−/−^* mice. Tau phosphorylation contributes to cognitive decline by disrupting microtubule stability, thereby leading to synaptic dysfunction, neuronal loss, and impaired memory formation. NF-κB signaling, which was downregulated in STAT4-deficient mice in NanoString analysis, has also been implicated in driving Tau phosphorylation and seeding throughout the brain (Wang et al., 2022). Additionally, GSEA of this NanoString data revealed that STAT4 deficiency resulted in a coordinated downregulation of the post-transcriptional modification pathway, which, in the context of improved memory and reduced Tau phosphorylation, could also indicate reduced Tau pathology spreading. In this pathway, genes involved in ubiquitination and subsequent degradation were downregulated with STAT4 deficiency. Tau is known to undergo ubiquitination following pathological phosphorylation, and this secondary modification may contribute to the formation of insoluble neurofibrillary tangles.(Li et al., 2021; Rawat et al., 2022; Yang et al., 2025). Recently, Zhang and colleagues reported that STAT4 deficiency under control of the LysM^Cre^-promotor improves long-term potentiation in the hippocampus of *Ldlr^−/−^* mice (Zhang et al., 2023). This is further supported by increased expression of the Rhodopsin-like signaling pathway in STAT4 deficiency, with *Cxcl16* specifically playing a neuroprotective role and modulating synaptic transmission in the CA1 of the hippocampus (Di Castro et al., 2016). Thus, the improved cognitive phenotype observed in *Stat4^ΔLysM^Ldlr^−/−^* vs *Stat4^fl/fl^Ldlr^−/−^* mice may involve enhanced synaptic plasticity.

Alternatively, the decrease in pTau burden could be due to amelioration of hypoperfusion. We have previously reported that LysM^Cre^-mediated STAT4 deficiency reduces atherosclerosis in the aortic sinus (Keeter et al., 2023). Stiffening of the aorta has been associated with damage to the microvasculature and consequent hypoperfusion of the brain (de la Torre, 2012). Therefore, by reducing atherosclerosis and aortic stiffness via STAT4 deficiency, cerebral blood flow may be preserved or improved, thereby reducing hypoperfusion-induced neuronal damage and pTau accumulation.

A detrimental effect of HFD on cognitive function has been documented in several reports, attributed to microgliosis, elevated levels of IL-1β, IL-6, and TNFα, and altered neuronal circuit function (Spencer et al.). Here, we show that LysM^Cre^-induced STAT4 deletion reduces the overall number of CD45^+^ leukocytes and microglia in the brain of *Stat4^ΔLysM^Ldlr^−/−^* mice, highlighting a role of STAT4 in the regulation of microgliosis and brain immunity. Multiple mechanisms may be responsible for altering brain immune composition, including a direct effect of STAT4 on the activation of LysM-targeted myeloid cells, and indirect effects on other cells, such as the activation of T and B cells, as evidenced by the expression of the early activation marker CD69. The LysM promoter efficiently targets myeloid lineage cells, including monocytes, macrophages, neutrophils, and dendritic cells (Mehrpouya-Bahrami et al., 2021). Additionally, Orthgiess and colleagues reported that LysM exhibits low-level activity in microglia and some neurons, but whether this activity affects the expression of floxed genes remains to be determined (Orthgiess et al., 2016).

Based on RNAseq data, STAT4 deletion leads to significant regulation of pathways associated with lipid metabolism and atherosclerosis, chemokine and cytokine signaling, and the AD-associated network. The role of STAT4 in inducing pro-inflammatory mediators, such as IFNγ, TNFα, and various chemokines that promote inflammation and activate the immune response, is well established. Here, we uncovered a pathological role of STAT4 in supporting neuroinflammation. Surprisingly, STAT4 deficiency had minimal effects on the number of neutrophils in the brain but reduced the proportion of immature CD64^+^ and CD115^+^ myeloid cells, suggesting that STAT4 may be involved in monocyte recruitment and maturation in the brain during atherogenesis. Interestingly, although STAT4 deletion was directed by LysM^Cre^ expression, RNAseq data revealed STAT4-dependent changes in BCR- and TCR-signaling pathways. These changes are likely driven by the overall alterations in the pro-inflammatory landscape of the brains of *Stat4^ΔLysM^Ldlr^−/−^* mice. Consistent with these observations, FACS analysis of the brains revealed upregulation of CD69 expression on T and B cells upon HFD-C feeding, which was reduced in the absence of STAT4. This finding provides further evidence of the role of STAT4 in regulating the adaptive immune response in the brains of HFD-C-fed atherosclerotic mice. We also observed increased viability in NeuN^+^ neurons with STAT4 deficiency. As STAT4 deficiency decreased several inflammatory pathways, it is possible that this improved neuronal viability was due to decreased neuroinflammation. Taupathology and hyperlipidemia-induced neuroinflammation are both known to drive increased neuronal death, both of which are implicated in this study (Bai et al., 2021; Tanabe et al., 2025). However, these changes in neuronal cell viability may also reflect improved clearance of dead cells via efferocytosis. Lipotoxicity due to overnutrition has been linked to impairments in efferocytosis, suggesting a potential dual cause of changes in neuronal viability that warrants further investigation (Mann et al., 2024).

The changes in the immune landscape of HFD-fed *Stat4^ΔLysM^Ldlr^−/−^* mice were reflected in efferocytosis assays. Efferocytosis in the brain, primarily mediated by microglia, plays a vital role in promoting cognitive health by efficiently clearing apoptotic cells and cellular debris. This process becomes especially critical in atherosclerosis, where chronic inflammation drives the activation and apoptosis of macrophages loaded with modified lipids. Interestingly, we found that the relative proportion, but not the number of efferocytosis-positive microglia and monocyte-derived macrophages, significantly increased in HFD-C-fed *Stat4^fl/fl^Ldlr^−/−^* vs chow diet-fed *Stat4^fl/fl^ Ldlr^−/−^* mice in *ex vivo* efferocytosis assays. Notably, STAT4 deficiency enhanced microglial efferocytosis, with a lesser effect on macrophage efferocytosis of apoptotic cells, suggesting that the STAT4/IL-12 axis regulates a key microglial function. STAT4 deficiency may facilitate a reparative shift in microglia by promoting the release of anti-inflammatory cytokines, such as IL-10 and TGFβ, following efferocytosis, thereby aiding the resolution of neuroinflammation and maintaining CNS homeostasis.

## Conclusion

In summary, several pathways driven by atherosclerosis could play a significant role in the development of neuroinflammation and associated cognitive decline. Atherosclerosis-related vascular dysfunction can impair small blood vessels, compromise the blood-brain barrier, trigger inflammation, and lead to neurovascular impairments. Systemic low-grade inflammation, reflected in the activation of both innate and adaptive immune arms, may support the recruitment of activated leukocytes to the brain. Additionally, hyperlipidemia associated with atherogenesis may alter the metabolic landscape, thereby impacting brain function. This study supports the critical role of STAT4 in shaping myeloid cell-driven inflammation, identifying STAT4 as a key molecular link between peripheral vascular inflammation and neuroinflammation that contributes to cognitive decline.

## Author contributions

Natalie Stahr: Writing - conceptualization, original draft, methodology, data curation, investigation. Alina Moriarty: data collection, data curation, formal analysis, writing. Shelby Ma: formal analysis, methodology, validation. Coles Keeter: data collection, analysis. Woong-Ki Kim and Larry Sanford: review and editing. Elena Galkina: conceptualization, supervision, funding, writing, review, and editing. All authors reviewed and approved the final draft before submission.

## Data availability statement

All data associated with this study are present in the paper. Data will be made available upon reasonable request.

